# A UG5 reverse transcriptase-nitrilase antiviral module confers phage immunity in the plant symbiont *Sinorhizobium meliloti*

**DOI:** 10.64898/2025.12.16.694600

**Authors:** Esperanza Sánchez-Nieto, Francisco Martínez-Abarca, Vicenta Millán, María Dolores Molina-Sánchez, Fernando M. García-Rodríguez, Nicolás Toro

## Abstract

Bacteriophages exert strong selective pressure on soil- and rhizosphere-associated bacteria, including plant-associated symbionts. Reverse transcriptase-associated defense systems of the UG family are widespread across bacterial lineages, yet their ecological roles remain largely undefined. Within this family, UG5 systems are distinguished by reverse transcriptases fused to or associated with a nitrilase domain. Here, we combine phylogenetic, metagenomic, and functional analyses to investigate the evolutionary context and antiviral activity of UG5-associated systems. Phylogenetic analysis of 728 nitrilase domains places UG5-associated nitrilases within a well-supported UG-related radiation encompassing the UG1, UG5, and UG6 families, with UG1 nested within a broader UG5 lineage. Metagenomic analysis further revealed UG5-associated reverse transcriptases in soil- and rhizosphere-derived metagenomes. Based on this observation, we characterized a UG5-large RT-associated system, here designated DRT11, encoded on the pSymA megaplasmid of *Sinorhizobium meliloti* RMO17, a nitrogen-fixing symbiont of *Medicago sativa*. Despite lacking the transmembrane protein typical of canonical UG5-large architectures, DRT11 confers protection against naturally occurring *M. sativa* rhizosphere phages with Podoviridae-like morphology. Phage infection assays reveal protection at low multiplicities of infection, consistent with an abortive-infection-like mechanism. Moreover, mutational analyses demonstrate that antiviral activity requires only the RT and its fused C-terminal nitrilase domain, establishing DRT11 as a minimal UG5-associated antiviral system.

## Introduction

Soil environments constitute one of the most complex microbial ecosystems on Earth, characterized by pronounced microscale heterogeneity and intense competition for limited resources. In these habitats, bacterial populations undergo continuous turnover driven by nutrient flow, protozoan grazing and, crucially, bacteriophage predation, which often represents a major source of microbial mortality (Emerson et al., 2018). Phages impose strong selective pressures that accelerate bacterial diversification through, lysogeny, genome erosion and the acquisition of horizontally transferred defense systems, thereby shaping microbial community structure (Touchon et al., 2017; Bernheim & Sorek, 2020). These forces are particularly pronounced in the rhizosphere, a biologically enriched compartment influenced by root exudates, where microbial densities and phage titers are high and ecological interactions strongly impact plant productivity (Philippot et al., 2013; Zhalnina et al., 2018; Wang et al., 2024). Yet, despite its ecological and agronomic relevance, the interplay between phage predation and bacterial immunity in soil-dwelling symbionts remains a largely unexplored frontier, with substantial implications for rhizosphere stability and for symbiotic interactions such as legume–rhizobium mutualism and the performance of nitrogen-fixing bioinoculants (Evans et al., 1979; Sharma et al., 2002).

*Sinorhizobium meliloti*, the nitrogen-fixing symbiont of alfalfa (*Medicago sativa*), must persist across both free-living and symbiotic stages to maintain an effective mutualism (Jones et al., 2007). Throughout this life cycle, rhizobial populations are continuously exposed to a diverse repertoire of lytic and temperate phages. Cultivation-based and metagenomic analyses show that *S. meliloti* phages exhibit substantial morphological and genomic diversity, including Siphoviridae, Myoviridae and Podoviridae members (Dziewit et al., 2014; Cubo et al., 2020). Together, these observations underscore that phage predation is a persistent and potent selective force in *S. meliloti*, likely contributing to the maintenance and diversification of antiviral defense systems. As phage-based approaches are increasingly developed for medical and agricultural applications (Jo et al., 2023; Villalpando-Aguilar et al., 2022; Hoffmann et al., 2025), mechanistic insight into antiviral pathways in beneficial soil bacteria will be essential to harness and protect these microbes in sustainable crop production.

Within this broader landscape of bacterial immunity, UG/Abi elements constitute a highly diverse family of prokaryotic reverse transcriptases (RTs) enriched in defense islands and widely predicted to mediate anti-phage activity (Mestre et al., 2022). Among the fortytwo UG/Abi RT groups described, UG5 comprises large class 3 RTs (typically >1000 aa) associated with mobile genetic elements and considerable structural variability. Although UG5-like modules with similar architectures have been implicated in antiviral activity, these systems were not formally classified as UG5 (Rousset et al., 2022), and bona fide UG5 representatives therefore remain functionally uncharacterized as defense systems.

In this study, we functionally analyzed a UG5-large system (WP_040120558.1) encoded on the megaplasmid pSymA of *Sinorhizobium meliloti* RMO17, a variant that notably lacks the transmembrane protein typically associated with UG5-large architectures (Mestre et al., 2022). We show that this system confers robust anti-phage immunity through a dual-component mechanism comprising only the RT and nitrilase domains, thereby expanding the known diversity of UG5-large systems and revealing a distinct antiviral activity in *S. meliloti*, where UG5 modules are rare and remain uncharacterized. Together, our findings provide one of the first molecular descriptions of a UG5-derived antiviral system in a plant-associated symbiont and open new avenues for understanding RT-linked defenses that protect rhizobial bacteria from phage predation, a feature that may be relevant in phage-enriched environments or in strategies combining bioinoculants with defined phage pressures.

## Materials and methods

### Bacterial strains, phages and growth conditions

*S. meliloti* strain RMO17 was originally isolated from root nodules of *Medicago orbicularis* growing in mildly acidic soils in Riego de la Vega (León, Spain) (Villadas et al., 1995). *S. meliloti* strain GR4 was isolated from nodules of *Medicago sativa* at the Estación Experimental del Zaidín field site (Granada, Spain) (Casadesús & Olivares, 1979). A bacteriophage collection infecting GR4 was obtained from *M. sativa* nodules collected in the la Vega region (Granada, Spain). This collection comprised 24 phages obtained from three independent isolation experiments, each designated using a two-part numerical code (1.1–1.11, 2.1–2.5 and 3.1–3.8) that denotes its isolate of origin (Toro et al., 2017). *S. meliloti* strains were cultivated in TY liquid medium or on TY agar plates at 30 °C. For plasmid mobilization into *S. meliloti*, *Escherichia coli* DH5α and HB101 were used as donor and helper strains, respectively, in triparental conjugation. DH5α (Bethesda Research Laboratories) and HB101 (Promega) were grown in LB liquid medium or on LB agar at 37 °C. When required, culture media were supplemented with the appropriate antibiotics to ensure plasmid maintenance.

### DNA oligonucleotides

DNA oligonucleotides used for conventional PCR, overlap extension PCR or sitedirected mutagenesis were synthesized by Metabion (Metabion International AG, Germany) and are listed in **Table S1**.

### Plasmid constructions

Plasmids constructed in-house by restriction enzyme cloning in this study are listed in **Table S2**. All constructs were verified by DNA sequencing. To express the full-length wild-type (WT) UG5-large system from *S. meliloti* RMO17 (WP_040120558.1), we constructed plasmid pBB2.1_UG5_WT. The expression vector used for this purpose was pBB2.1, a kanamycin-resistant (Km^R^) derivative of pBB2 in which the native SalI site was removed to facilitate SalI-compatible cloning steps. The UG5 coding sequence (CDS) was first amplified by PCR (see section “UG5-based PCR”) and cloned into the ampicillin-resistant (Ap^R^) pSpark® I vector (Canvax), following the manufacturer’s instructions, generating pSparkI_UG5_WT. This plasmid was digested with HindIII and SpeI to release the UG5 insert, which was ligated into HindIII/SpeI-digested pBB2, yielding pBB2_UG5_WT. To remove a 179 bp fragment located between two SmaI restriction sites, pBB2_UG5_WT was digested with SmaI, generating pBB2_UG5_Δ179. The modified UG5 insert was then excised with HindIII and SpeI and ligated into HindIII/SpeI-digested pBB2.1, generating the final construct pBB2.1_UG5_WT used in all subsequent cloning steps.

To generate the mutant carrying the DD296–297AA substitution in the RT domain, we constructed plasmid pBB2.1_UG5_RT[DD296–297AA]. A three-step PCR strategy was used to produce two overlapping fragments containing the desired mutation, which were subsequently fused by overlap extension PCR (see section “RT mutant fragment-based overlap extension PCR” for details). The resulting full-length amplicon was digested with BamHI and SalI and ligated into BamHI/SalI-digested pBB2.1_UG5_WT, replacing the corresponding wild-type region and yielding the final construct pBB2.1_UG5_RT[DD296–297AA].

To generate the mutant carrying the C876A substitution in the nitrilase domain, we constructed plasmid pBB2.1_UG5_Nitrilase[C876A]. A synthetic DNA fragment encompassing the 3′ region of UG5 (between the SalI and SpeI restriction sites) and incorporating the desired point mutation was obtained from GenScript Biotech. The fragment, delivered in an Ap^R^ pUC57m vector, was excised with SalI and SpeI and ligated into SalI/SpeI-digested pBB2.1_UG5_WT, thereby replacing the corresponding wildtype region and yielding the final construct pBB2.1_UG5_Nitrilase[C876A].

### Bacterial total DNA isolation

Bacterial total DNA was extracted from 5 mL cultures of *S. meliloti* RMO17 using the REAL Genomic DNA Bacteria kit (REAL Laboratory) following the manufacturer’s instructions, with an initial lysozyme digestion step to facilitate lysis. Briefly, cell pellets collected at an OD₆₀₀ of 0.5 were resuspended in lysis buffer supplemented with lysozyme (10 mg mL⁻¹) and incubated for 1 h at 37 °C. After lysis, samples were treated with RNase, subjected to protein precipitation, and genomic DNA was recovered by isopropanol precipitation and washed with 70% ethanol. Pellets were air-dried and resuspended in Milli-Q water, followed by incubation at 65 °C to facilitate hydration. DNA concentration was quantified using a Qubit® 2.0 fluorometer (Invitrogen), and samples were stored at –20 °C until further use.

### UG5-based PCR

PCR amplification of the full-length UG5 locus (WP_040120558.1) from *S. meliloti* RMO17 genomic DNA was performed using primers UG5_Fw and UG5_Rv (**Table S1**). Reactions (50 µL) contained 0.1–100 ng template DNA, 200 µM dNTPs, 500 nM of each primer, 1× Phusion HF buffer and 0.5 µL Phusion High-Fidelity DNA Polymerase (Thermo Scientific). Amplification was carried out in a Mastercycler® Nexus gradient thermocycler (Eppendorf) with the following program: 98 °C for 30 s; 30 cycles of 98 °C for 20 s, 60 °C for 30 s, and 72 °C for 2 min; and a final extension at 72 °C for 5 min. Negative controls lacking template DNA were included in all experiments. PCR products were examined by agarose gel electrophoresis.

### RT mutant fragment-based overlap extension PCR

Two partially overlapping UG5 fragments were amplified from plasmid pBB2.1_UG5_WT using primer pairs 2626_Fw / YVAA_Rv and YVAA_Fw / 4370_Rv (**Table S1**). The resulting products were fused by overlap extension PCR with the outer primers 2626_Fw and 4370_Rv. Reactions (50 µL) contained 0.1–100 ng template DNA, 200 µM dNTPs, 500 nM of each primer, 1× Phusion HF buffer and 0.5 µL Phusion High-Fidelity DNA Polymerase (Thermo Scientific). Amplifications were performed in a Mastercycler® Nexus gradient thermocycler (Eppendorf). For PCR A and PCR B, cycling consisted of 98 °C for 30 s; 25 cycles of 98 °C for 20 s, 62 °C for 20 s, and 72 °C for 45 s; followed by 72 °C for 5 min. The overlap-extension reaction (PCR C) used identical conditions except for an annealing temperature of 60 °C and an extension of 1 min. Negative controls lacking template DNA were included throughout. Final products were assessed by agarose gel electrophoresis.

### Large terminase (TerL)-based PCR

PCR amplification of the *terL* gene from phages 1.3 and 3.2 was performed using the corresponding phage stocks as DNA templates and the primer pairs 1.3TerL-881_Fw / 1.3TerL-1558_Rv and 3.2TerL-807_Fw / 3.2TerL-1261_Rv (**Table S1**). Reactions (20 µL) contained 2 µL of phage stock, 200 µM dNTPs, 500 nM of each primer, 1× reaction buffer (50 mM KCl, 2.5 mM MgCl₂, 10 mM Tris-HCl, pH 8.3), 0.1 µL of homemade Taq polymerase, and Milli-Q® water to volume. Amplifications were performed in a Mastercycler® nexus gradient thermocycler (Eppendorf) with an initial denaturation at 94 °C for 30 s, followed by 30 cycles of 94 °C for 20 s, 60 °C for 30 s, and 72 °C for 30 s, and a final extension at 72 °C for 5 min. A no-template control was included in each set of reactions. PCR products were resolved by non-denaturing agarose gel electrophoresis.

### Transformation of *E. coli* DH5*α* by heat shock

Chemically competent *E. coli* DH5α cells were prepared using the rubidium chloride method of Rodríguez & Tait (1983). Aliquots of 100 µL were thawed on ice for ∼20 min, after which 50–500 ng of plasmid DNA was added and incubated on ice for 30 min. Cells were heat-shocked at 42 °C for 2 min and immediately returned to ice for 5 min. For recovery, 900 µL of LB medium was added, and cultures were incubated at 37 °C for 1 h with gentle agitation. Cells were then plated onto LB agar containing the appropriate antibiotic and incubated overnight at 37 °C. The plated volume was adjusted according to the expected transformation efficiency and DNA input.

### Plasmid DNA isolation from *E. coli*

Plasmid DNA was isolated from 1.5 mL *E. coli* cultures using a magnesium-salt precipitation method as described by Studier (1991). Overnight cultures were pelleted by centrifugation at 13,000 rpm for 3 min at room temperature, resuspended in 100 µL Milli-Q® water, and lysed by addition of 100 µL lysis buffer (10 mM EDTA, 0.1 M NaOH, 2% SDS) with immediate mixing. Samples were incubated for 2 min in a boiling-water bath. Lysis was quenched by adding 50 µL of 1 M MgCl₂, vortexing briefly, and placing the tubes on ice for 5 min to precipitate linear chromosomal DNA. After centrifugation (13,000 rpm, 1 min), 50 µL of 5 M potassium acetate (pH 4.8) were added to the supernatants to remove residual proteins. Following incubation on ice for 2 min and centrifugation (13,000 rpm, 5 min), the cleared supernatants were transferred to fresh tubes containing 600 µL cold 100% ethanol. DNA was precipitated on ice for 5 min and pelleted by centrifugation at 13,000 rpm for 5 min. Pellets were washed with 200 µL cold 70% ethanol, dried (air-dry or 10–15 min in a SpeedVac), and resuspended in 10–30 µL of RNase-containing Milli-Q® water (10 µg µL⁻¹ final). Plasmid DNA was stored at –20 °C until use.

### Plasmid transfer to *S. meliloti* GR4 by triparental conjugation

Plasmid mobilization into *S. meliloti* GR4 was performed by triparental conjugation (Ditta et al., 1980) using *E. coli* DH5α as the donor strain and *E. coli* HB101 carrying the helper plasmid pRK2013. Biomass from overnight cultures of the three strains was scraped from TY agar plates, mixed in approximately equal proportions, and spotted onto fresh TY agar. After overnight incubation at 30 °C, the mixture was streaked onto MM agar containing the appropriate antibiotic to select for *S. meliloti* transconjugants while counter selecting *E. coli*. Plates were incubated for 2–3 days at 30 °C until isolated colonies were obtained.

To verify strain identity, individual colonies were replica-plated onto fresh MM agar and, in parallel, onto Endo agar. MM plates were incubated overnight at 30 °C, whereas Endo agar plates were incubated overnight at 37 °C. Colonies that grew on MM agar but failed to grow on Endo agar were scored as *S. meliloti* GR4 transconjugants.

### Plasmid DNA isolation from *S. meliloti* by alkaline lysis

Plasmid DNA was isolated from 1.5 mL of *S. meliloti* cultures by alkaline lysis (Sambrook et al., 1989) based on the methods of Birnboim and Doly (1979) and Ish-Horowicz and Burke (1981). Overnight cultures were pelleted by centrifugation at 13,000 rpm for 3 min at room temperature, washed with 500 µL of 0.1% sarcosyl in TE buffer, and centrifuged again under the same conditions. Pellets were resuspended in 100 µL of lysozyme (4 mg mL⁻¹) prepared in solution I (10 mM EDTA, 50 mM glucose, 25 mM Tris-HCl pH 8.0) and incubated for 5 min at room temperature. Cell lysis was induced by adding 200 µL of solution II (0.2 M NaOH, 1% SDS) and gently inverting the tubes, followed by incubation on ice for 5 min. Neutralization was achieved by adding 150 µL of 5 M potassium acetate (pH 4.8), mixing, and incubating on ice for 5 min. Samples were centrifuged at 13,000 rpm for 5 min, and the supernatants were transferred to fresh tubes.

To remove proteins, the lysates were extracted with an equal volume of phenol:chloroform:isoamyl alcohol (25:24:1), vortexed, and centrifuged at 13,000 rpm for 5 min. The aqueous phase was recovered and further extracted with chloroform:isoamyl alcohol (24:1). Plasmid DNA was precipitated by adding 2.5 volumes of 100% cold ethanol and incubating for 20 min at –20 °C (or 15 min at –80 °C). After centrifugation at 13,000 rpm for 5 min, pellets were washed with 70% cold ethanol, centrifuged, air-dried or SpeedVac-dried, and resuspended in 10–30 µL of RNase-containing Milli-Q water (10 µg µL⁻¹ RNase from a 100× stock). Plasmid DNA was stored at –20 °C.

### Phage plaque assays

Double-layer plaque assays were performed following Kropinski et al. (2009). *S. meliloti* GR4 strains were grown overnight at 30 °C in TY medium supplemented with the appropriate antibiotic. Bacterial lawns were prepared by mixing 300 µL of culture with 10 mL of molten top TY agar (0.5% agar, 5 mM MgSO₄, plus antibiotic), which was poured onto square plates (12 × 12 cm) containing bottom TY agar (1.5% agar, 5 mM MgSO₄, plus antibiotic). Serial 10-fold phage dilutions were prepared in phage buffer (TY supplemented with 5 mM MgSO₄), and 10 µL of each dilution was spotted onto the solidified lawns. Plates were incubated at 30 °C for 24–48 h, after which plaques were counted and phage titers were calculated as PFU mL⁻¹. When plaques were too small to be quantified reliably, the most concentrated dilution lacking visible plaques was recorded as corresponding to a single plaque.

### Phage infection assays in liquid cultures

Phage-infection assays in liquid culture were performed essentially as described by Gapińska et al. (2024), with minor modifications. Pre-inocula were prepared by streaking three independent colonies of *S. meliloti* GR4 strains carrying either the empty-vector control or the UG5 WT system into 3 mL TY medium supplemented with the appropriate antibiotic, followed by incubation for 2 days at 30 °C. Saturated cultures were diluted 1:1000 into 4 mL of fresh TY medium with antibiotic and grown overnight at 30 °C. The following morning, OD₆₀₀ was measured and cultures were adjusted to an OD₆₀₀ of 0.2. For infection assays, 180 µL of each normalized culture were dispensed into wells of a 96-well plate, and 20 µL of freshly prepared phage suspensions were added to achieve final multiplicities of infection (MOI) of 0 (no phage), 0.02, 0.2, and 2. Control wells received 20 µL of TY medium. Bacterial growth was monitored at 30 °C in a Varioskan™ LUX plate reader (Thermo Scientific), with OD₆₀₀ measurements taken every 30 min for 17 h.

### Transmission electron microscopy

Phage were examined by transmission electron microscopy at the Confocal and Microscopy Service (CTEM) of the Estación Experimental del Zaidín (CSIC; Granada, Spain), following standard negative staining procedures with uranyl acetate.

### Phage total DNA isolation

Phage total DNA was extracted from 1 mL of high-titer lysates (≥10⁹ PFU mL⁻¹) or from 1.5 mL when titers were lower. Lysates were mixed with 0.2 volumes of M1 buffer (3 M NaCl, 30% polyethylene glycol 6000), vortexed briefly, incubated on ice for 1 h, and centrifuged at 13,000 rpm for 10 min at room temperature to pellet phage particles. Supernatants were discarded, and pellets were resuspended in 200 µL of M2 buffer (500 mM guanidine-HCl, 10 mM MOPS, 1% Triton X-100; pH 6.5), incubated for 10 min at 80 °C to lyse virions, and cooled to room temperature for 10 min. Lysates were then mixed with 200 µL of MX3 buffer from the mi-Plasmid Miniprep kit (Metabion) to neutralize the solution and centrifuged at 13,000 rpm for 10 min. The resulting supernatants were transferred to Mini Plus columns from the same kit and centrifuged at 9,000 rpm for 1 min. Columns were washed sequentially with 500 µL of WN buffer and 700 µL of WS buffer, each followed by centrifugation at 9,000 rpm for 1 min. To remove residual ethanol, columns were centrifuged at 13,000 rpm for 3 min and transferred to clean tubes. Phage DNA was eluted with 30 µL of Milli-Q® water by centrifugation at 13,000 rpm for 3 min. DNA concentration was measured with a Qubit® 2.0 fluorometer (Invitrogen), and samples were stored at –20 °C until use.

### Phage genome sequencing and assembly

Phage DNA was sequenced at the Sequencing Service of Fisabio (Valencia, Spain) using the Illumina platform. Quality-checked and trimmed reads were assembled using Geneious Prime software (version 2020.0.2).

### Phylogenetic analyses

Tree reconstruction was performed with IQ-TREE2 (Minh et al., 2020) using the best-fit model identified by ModelFinder (Q.pfam+R10). Branch support values correspond to UFBoot2 and SH-aLRT, each estimated from 1,000 replicates.

## Results

### Gene architectures, phylogenetic structure and metagenomic distribution of UG5-associated nitrilases

Two alternative architectures have been described for UG5 systems (Mestre et al., 2022). In UG5-large, the reverse transcriptase (RT) is C-terminally fused to a nitrilase domain and is often accompanied by a transmembrane (TM) protein from clusters 85 or 1858. By contrast, UG5-small encodes the nitrilase domain (cluster 249) in a separate open reading frame and lacks an associated TM protein. Beyond UG5, RTs from UG1, UG5 and UG29 also carry a C-terminal nitrilase domain, whereas in UG6 the nitrilase is encoded by an independent downstream gene (cluster 285).

Phylogenetic analysis of 728 nitrilase-domain sequences revealed a well resolved and deeply structured topology among UG-associated nitrilases within the broader nitrilase superfamily. The UG29-associated nitrilases form a strongly supported and clearly monophyletic clade (UFBoot = 100). In contrast, neither UG5-associated nor UG1-associated nitrilases are recovered as exclusive monophyletic groups. Instead, UG1 members are nested within a broader UG5-associated radiation: the least common ancestor defining the UG5 cluster is highly supported (UFBoot = 99) and encompasses all UG1 sequences, indicating that UG1 does not represent an independent lineage in this dataset. Nitrilases from CL249 and CL285, associated with UG5-small and UG6, respectively, also fall within this radiation and similarly fail to form strictly monophyletic groups.

The internal node uniting the UG1/UG5/CL249/CL285 assemblage is strongly supported (UFBoot ≈ 99), whereas the next higher node connecting this cluster to the remainder of the superfamily shows only moderate support (UFBoot ≈ 73). Collectively, these results indicate that nitrilases associated with UG5, UG1, CL249 and CL285 constitute a single, well-supported evolutionary group whose diversification does not partition into discrete monophyletic families. In contrast, UG29 branches as a distinct and robustly supported lineage that is clearly separated from the UG1/UG5-associated radiation (**Figure 1A, Data file 1**).

**Figure 1.**
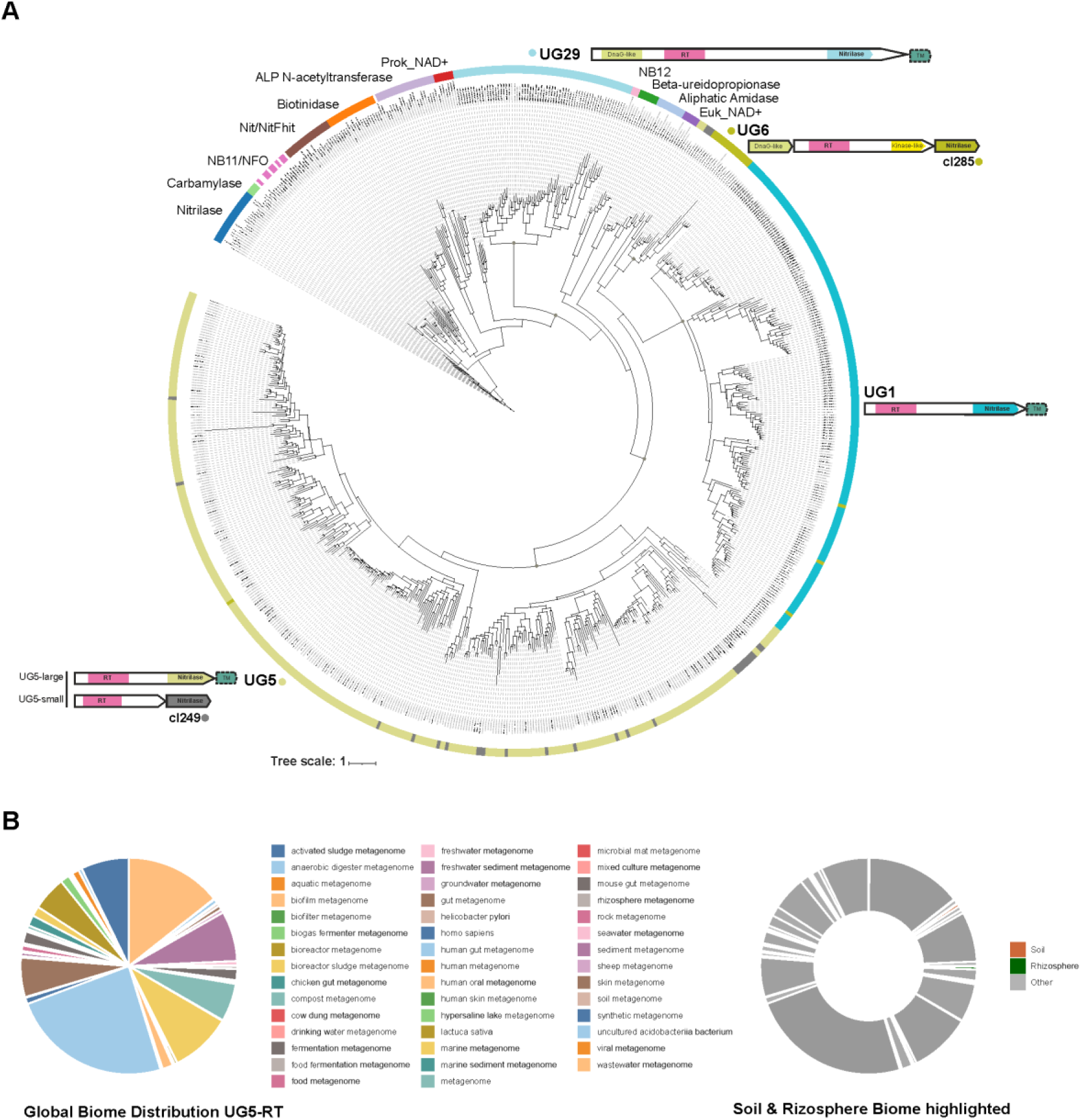
Gene architecture, phylogenetic relationships and metagenomic distribution of UG5 systems. (**A**) Phylogenetic analysis of nitrilase domains across UG retron-associated systems and canonical CN-hydrolase family members. A maximum-likelihood phylogeny was inferred from a curated alignment of 728 nitrilase-domain sequences, including the UG5-large nitrilase, the CL249 nitrilase associated with UG5-small, the nitrilases linked to UG1 and UG29, and the CL285 nitrilase associated with UG6, together with representative members of the broader nitrilase superfamily (Pace and Brenner, 2001). The phylogeny is shown as a circular tree, with outer colored ring segments denoting major clades. Key internal nodes resolving relationships among UG/Abi-associated nitrilases are highlighted with filled circles (UFBoot ≥ 95%). Schematic domain architectures of UG5-, UG1-, UG6- and UG29-associated nitrilases are displayed in the outer panel to contextualize their phylogenetic placement. (**B**) Metagenomic distribution of UG5-associated reverse transcriptase domain across environmental biomes. UG5-associated reverse transcriptase (RT) sequences were identified by HMM-based searches against the MGnify protein database using a profile derived from the UG5 RT domain. Retrieved hits were filtered using a stringent significance threshold (E-value ≤ 1 × 10⁻⁵⁰) to retain high-confidence sequences, which were subsequently mapped to sample-level biome annotations. The figure shows the global distribution of UG5-associated RT sequences across annotated biomes, together with a highlighted hybrid donut representation in which soil- and rhizosphere-derived metagenomes are shown explicitly and all remaining biomes are grouped as “Other.” The overall biome distribution closely mirrors that obtained using the UG5 nitrilase-domain (**Figure S1**) HMM, supporting the robustness of the metagenomic analysis.

To place this evolutionary framework in a broader ecological context, we next examined the environmental distribution of UG5-related systems using a metagenomic approach. An HMM derived from the UG5 reverse transcriptase domain was used to query the MGnify protein database, yielding more than 27,000 initial matches across a wide range of metagenomic datasets. Applying a stringent significance threshold (E-value ≤ 1 × 10⁻⁵⁰) reduced this dataset to 949 high-confidence RT matches, consistent with bona fide UG5-related systems. Mapping high-confidence UG5-RT hits to sample-level metagenomic biomes revealed a heterogeneous environmental distribution. UG5-associated RTs were identified across multiple metagenomic contexts, consistent with the broad environmental scope of the datasets analyzed. Soil- and rhizosphere-derived metagenomes together accounted for a relatively small fraction of filtered UG5-RT matches (approximately 0.84%), a value that should be interpreted cautiously given the annotation biases of large-scale metagenomic repositories (**Figure 1B).** Consistent results were obtained using an independent HMM search based on UG5-associated nitrilase sequences (**Figure S1**). Rather than indicating enrichment in a specific biome, this pattern highlights the recurrent presence of UG5-associated systems across diverse environments and provides contextual support for their functional investigation in soil- and rhizosphere-associated bacteria.

### Genomic and structural characterization of the UG5-large locus from *S. meliloti* RMO17

Given the ecological importance of antiviral defenses in soil-dwelling symbionts, and the fact that bona fide UG5 systems remain functionally unvalidated as defense modules, we aimed to characterize a representative UG5 locus in a biologically relevant nitrogen-fixing bacterium. To this end, we focused on the UG5-large variant (WP_040120558.1) encoded on the megaplasmid pSymA (CP009145.1) of *S. meliloti* RMO17 (genome assembly GCF_000747295.1), a strain whose genome was previously sequenced and is available in our laboratory.

The RMO17 genome comprises a chromosome (CP009144.1; 3,649,532 bp) and two megaplasmids, pSymB (CP009146.1; 1,610,737 bp) and pSymA (CP009145.1; 1,466,845 bp), as reported previously (Toro et al., 2014). The UG5-large locus, spanning coordinates 109,344–112,361 on pSymA, encodes a single 1005-aa polypeptide comprising an RT domain fused to a C-terminal nitrilase domain and notably lacks an associated TM protein (**Figure 2A**). Phylogenetically, this locus is most closely related to the UG5-2598 entry (WP_180162318.1) from *S. medicae* (assembly GCF_901933175.1) (Mestre *et al*., 2022), and homologous loci are also present on pSymA of other *S. meliloti* strains, including Rm41 (CP021811) and KH35C (CP021826), as well as in *Ensifer adhaerens* strains such as OV14 (CP007239).

**Figure 2.**
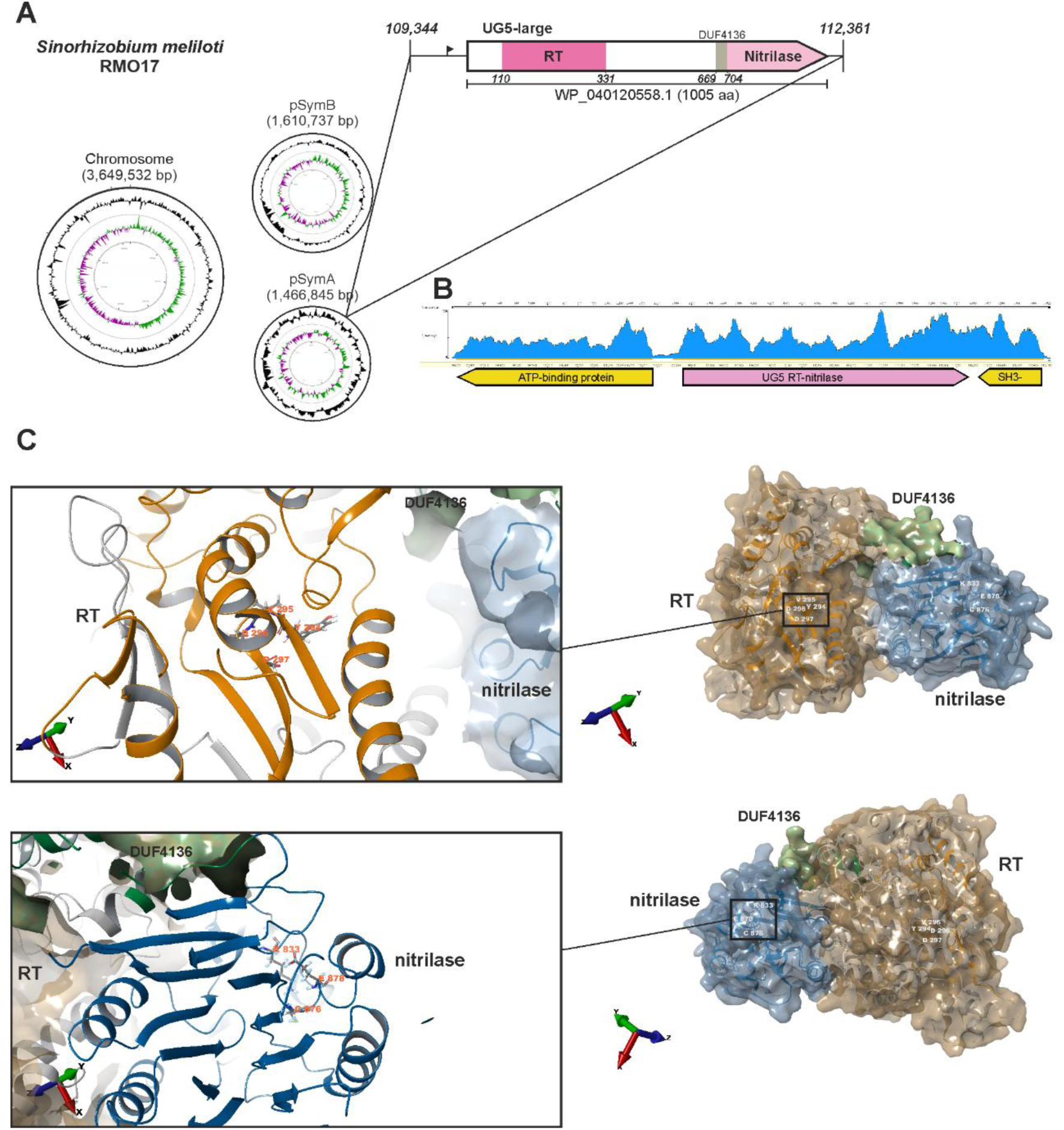
Genomic context and structural organization of the UG5-large system in *S. meliloti* RMO17. (**A**) Genomic organization of the UG5-large locus. The genome of RMO17 (assembly GCF_000747295.1) comprises a chromosome (CP009144.1; 3,649,532 bp) and two megaplasmids: pSymB (CP009146.1; 1,610,737 bp) and pSymA (CP009145.1; 1,466,845 bp). The UG5-large system (WP_040120558.1; 1005 aa) is encoded on pSymA, spanning coordinates 109,344–112,361, and consists of a reverse transcriptase (RT) fused to a C-terminal nitrilase domain. Notably, this locus lacks a co-located transmembrane protein. The arrow represents the predicted ORF, scaled to gene length and oriented according to transcriptional direction. Domain annotations are shown within the ORF. Colors denote domain identity (RT, bright pink; nitrilase, light pink). (**B**) Transcriptomic profiling of the UG5-large region in exponential growth phase *S. meliloti* RMO17. Illumina RNA-seq reads were mapped to the pSymA region encompassing the UG5-large locus, showing the coverage profile across the RT–nitrilase ORF and its neighboring genes. No discrete accumulation of reads was detected upstream or downstream of the UG5-large coding sequence under the conditions tested, and thus no transcript consistent with a co-localized ncRNA was observed. Annotated ORFs flanking the locus are shown below the coverage track. (**C**) AlphaFold3 (Abramson et al., 2024) structural model of the UG5 RT–nitrilase fusion shown in two orientations, displayed as a molecular surface and colored by domain. The protein adopts a modular architecture comprising an N-terminal RT domain, a small DUF4136 domain, and a C-terminal nitrilase (CN-hydrolase) domain. Ribbon traces are shown beneath the surface for clarity. Domain boundaries follow MyCLADE annotations (Vicedomini *et al*., 2021) and structural homology inferred from US-align (Zhang et al., 2022) comparisons with cluster 249 nitrilases (**Figure S2**). Insets highlight the catalytic centers of the RT (YVDD) and nitrilase (K–C–E) domains.

In class 3 UG/Abi systems, a ncRNA that could serve as a template for reverse transcription by the associated RT has not yet been identified. Consistent with this, transcriptomic analyses of *S. meliloti* RMO17 did not reveal any obvious accumulation of reads flanking the UG5 RT–nitrilase gene (**Figure 2B**), providing no support for the presence of a candidate ncRNA in the UG5-large system of this bacterium. The absence of a detectable ncRNA suggests that this UG5-large system may employ a protein-primed mode of synthesis, although further work will be required to distinguish this possibility from alternative mechanisms.

To gain structural insight into how the RT and nitrilase domains are organized within a single polypeptide, we generated a high-confidence AlphaFold3 model of the *S. meliloti* RMO17 encoded UG5-large protein (**Figure 2C**). The model reveals a compact RT fold followed by a smaller DUF4136 domain, which acts as a structural connector between the RT and the C-terminal nitrilase domain. Consistent with its assignment as a reverse transcriptase, the RT domain contains the conserved YVDD catalytic tetrad (residues 294–297), positioned in a canonical palm subdomain configuration. Likewise, the nitrilase domain displays the characteristic K–C–E catalytic triad (Lys833–Cys876– Glu878), arranged identically to bona fide CN-hydrolases.

### Genomic context of the RMO17 UG5-large system

To identify co-localized defense systems, a PADLOC-based analysis (Payne et al., 2022) was performed on the *S. meliloti* RMO17 genome, which revealed three additional putative defense systems adjacent to the UG5-large locus: (i) a nucleotidyl transferase paired with an AbiEii/AbiGii-like toxin (WP_020479331.1), (ii) a type II toxin-antitoxin system belonging to the VapC family (WP_040120557.1), and (iii) a type II toxin-antitoxin system comprising a Phd/YefM family antitoxin (WP_014528082.1). The UG5-large gene resides within a variable genomic region flanked by conserved sequences. To assess the distribution of this region across related strains, we performed a BLAST-based comparative analysis using the conserved flanks as queries. This analysis revealed that the variable region is present on the pSymA megaplasmid of other *S. meliloti* strains, including RCAM1750 (CP050513.1) and KH46 (CP021824.1), although the UG5-large system itself is absent. Despite this variability, the surrounding genomic context is highly conserved across strains (**Figure 3**). This pattern, conserved flanking sequences adjacent to a variable gene block present in only a subset of strains, is consistent with the horizontal acquisition of this region.

**Figure 3.**
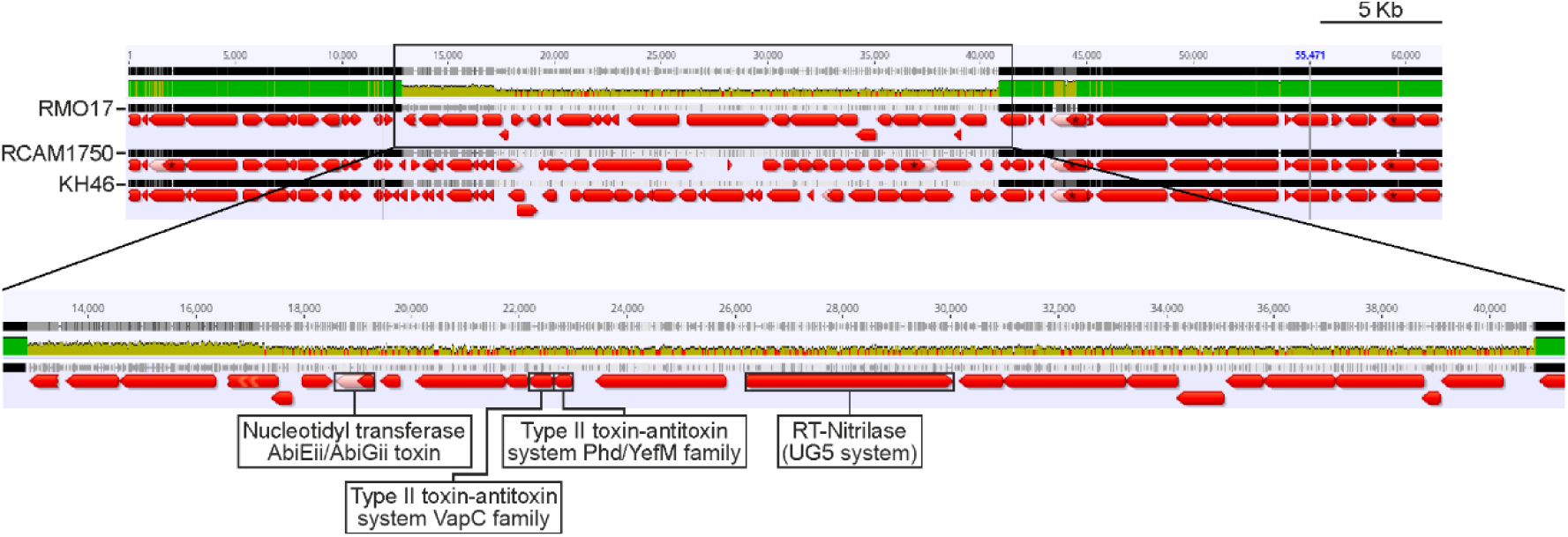
UG5-large in a variable genomic region. Comparative genomic context within the megaplasmid pSymA of *S. meliloti* strains RMO17 (CP009145.1), RCAM1750 (CP050513.1) and KH46 (CP021824.1), highlighting a shared variable region present in all three strains. Within this region, the UG5-large system (RT–nitrilase; WP_040120558.1) is found exclusively in RMO17. Adjacent to UG5-large, three additional putative defense systems were identified: a predicted nucleotidyl transferase paired with an AbiEii/AbiGii-like toxin (WP_020479331.1), a type II toxin–antitoxin module of the VapC family (WP_040120557.1), and a type II toxin–antitoxin module comprising a Phd/YefM family antitoxin (WP_014528082.1). The surrounding genomic context shows high nucleotide-level conservation across strains. Bright green blocks denote conserved regions, khaki blocks indicate variable segments, and red arrows represent predicted ORFs, scaled to gene length and oriented according to transcriptional direction.

### Anti-phage defense activity of the UG5-large system

To evaluate the defensive capacity of the UG5-large system, we expressed this locus in *S. meliloti* GR4, which naturally lacks it, and challenged the strain with a panel of 24 phages isolated from root nodules of *M. sativa* plants that were originally identified based on their ability to infect this host (Toro et al., 2017). Expression of the UG5-large system conferred robust protection, as evidenced by a marked reduction in efficiency of plating (EOP) observed for phages 1.4, 1.6, 2.1, 2.2 and 2.4–3.7 (**Figure 4A**).

**Figure 4.**
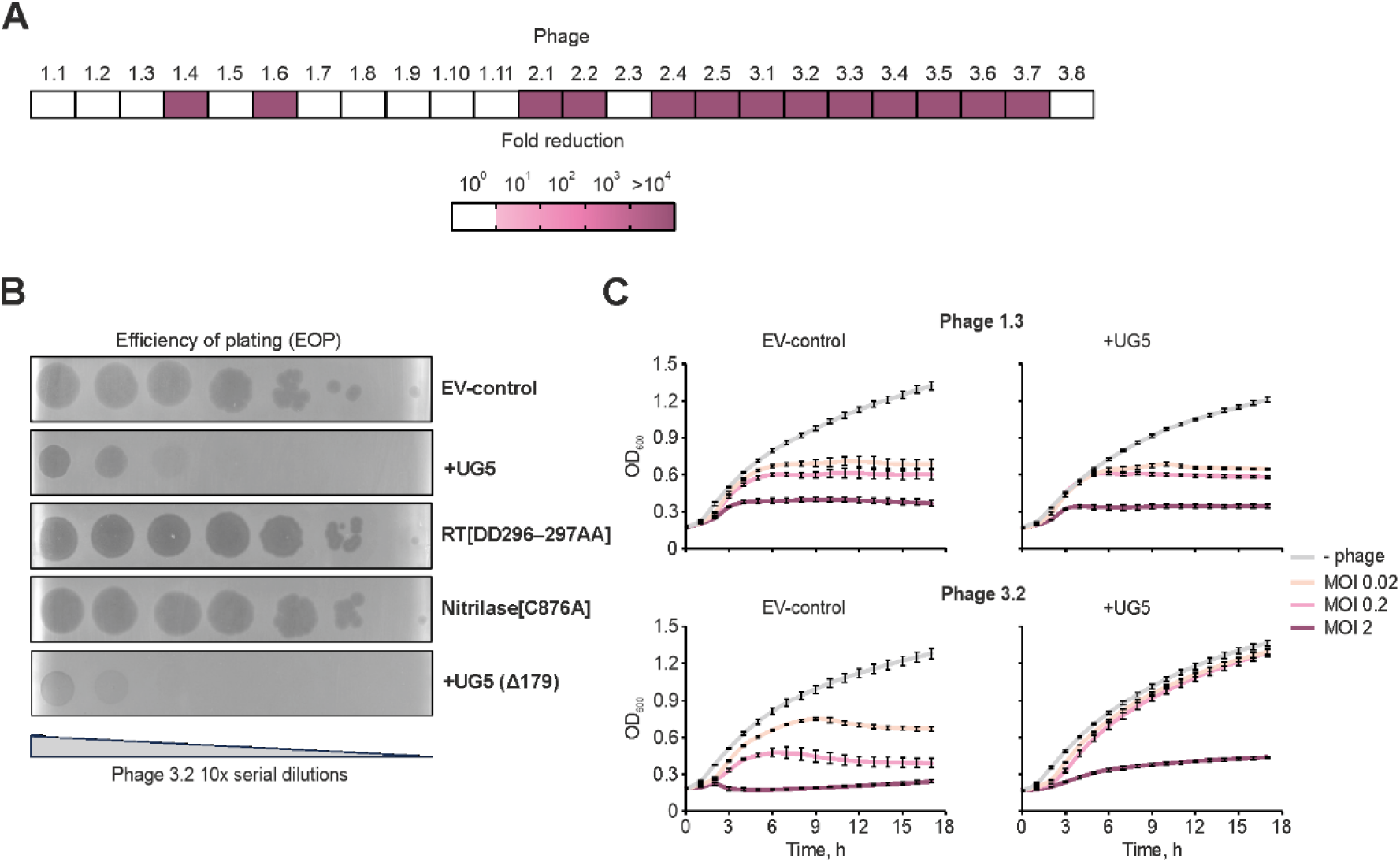
UG5-large-mediated phage resistance. **(A)** Heat map summarizing the fold reduction in phage infectivity conferred by the UG5-large system against 24 *S. meliloti* GR4 phages isolated from *Medicago sativa* root nodules collected in the Vega area (Granada, Spain). Fold reduction was determined by plaque assays using serial 10-fold dilutions, comparing the efficiency of plating (EOP) between a *S. meliloti* GR4 strain carrying the empty-vector (EV) control or the UG5-large system. **(B)** Serial 10-fold dilution plaque assays comparing the EOP of phage 3.2 on *S. meliloti* GR4 strains carrying the empty-vector control, the UG5 system, UG5 (Δ-179) carrying a deletion of 179 bp upstream the RT-nitrilase predicted promoter (positions 109,019 to 109,193) or catalytic-site mutants (RT[DD296–297AA] and Nitrilase[C876A]) expressed from the pBB2.1 vector. For clarity, only plaques corresponding to phage 3.2 are shown. Results are representative of three independent experiments. **(C)** Growth curves of *S. meliloti* GR4 strains carrying either the vector-only control or the UG5 system upon infection with phages 1.3 or 3.2 at multiplicities of infection (MOI) of 0, 0.02, 0.2, and 2. The UG5 system does not protect against phage 1.3, but confers MOI-dependent protection against phage 3.2, consistent with an abortive-infection mechanism. Bacterial growth was monitored over 17 h at 30 °C. Curves represent the mean of three biological replicates; error bars indicate the standard error of the mean.

To determine the functional contribution of each domain, we constructed catalytic mutants in the reverse transcriptase (RT[DD296–297AA]) and the nitrilase domain (Nitrilase[C876A]). Serial 10-fold dilution plaque assays with phage 3.2 showed that both mutations abolished UG5-large–mediated protection, yielding EOP values comparable to those of the vector-only control. Moreover, deletion of a 179-bp region upstream of the predicted RT–nitrilase promoter (positions 109,019–109,193) did not impair UG5 defense activity, further supporting the absence of a ncRNA template in this system (**Figure 4B**).

To further evaluate UG5-large defense activity, we performed phage infection assays in liquid culture at multiplicities of infection (MOI) of 0, 0.02, 0.2, and 2 using two representative phages: phage 1.3, which UG5-large does not protect against, and phage 3.2, which is effectively restricted by the system. In the absence of phage (MOI 0), the control and UG5-large strains exhibited comparable growth. Upon infection with phage 1.3, no differences were observed between strains across all MOIs. By contrast, phage 3.2 impaired growth of the control strain at all MOIs, whereas the UG5-large–expressing strain sustained growth at low MOI but collapsed at high MOI (**Figure 4C**). This MOI-dependent protection is characteristic of abortive infection systems, suggesting that UG5-large confers antiviral defense through an Abi-like mechanism. Given its experimentally validated activity, we hereafter refer to this UG5-large module as DRT11.

### Morphological and genomic characterization of GR4 phages

Morphological features of phages 1.3 and 3.2, which are respectively resistant and susceptible to DRT11-mediated defense, were examined by transmission electron microscopy at the Microscopy Service of the Estación Experimental del Zaidín–CSIC (Granada, Spain). As shown in **Figure 5A**, both phages display traits characteristic of the *Podoviridae* family, including an icosahedral capsid (approximately 50 nm in diameter) and a short, non-contractile tail (Cubo et al., 2020; Kornienko et al., 2020).

**Figure 5.**
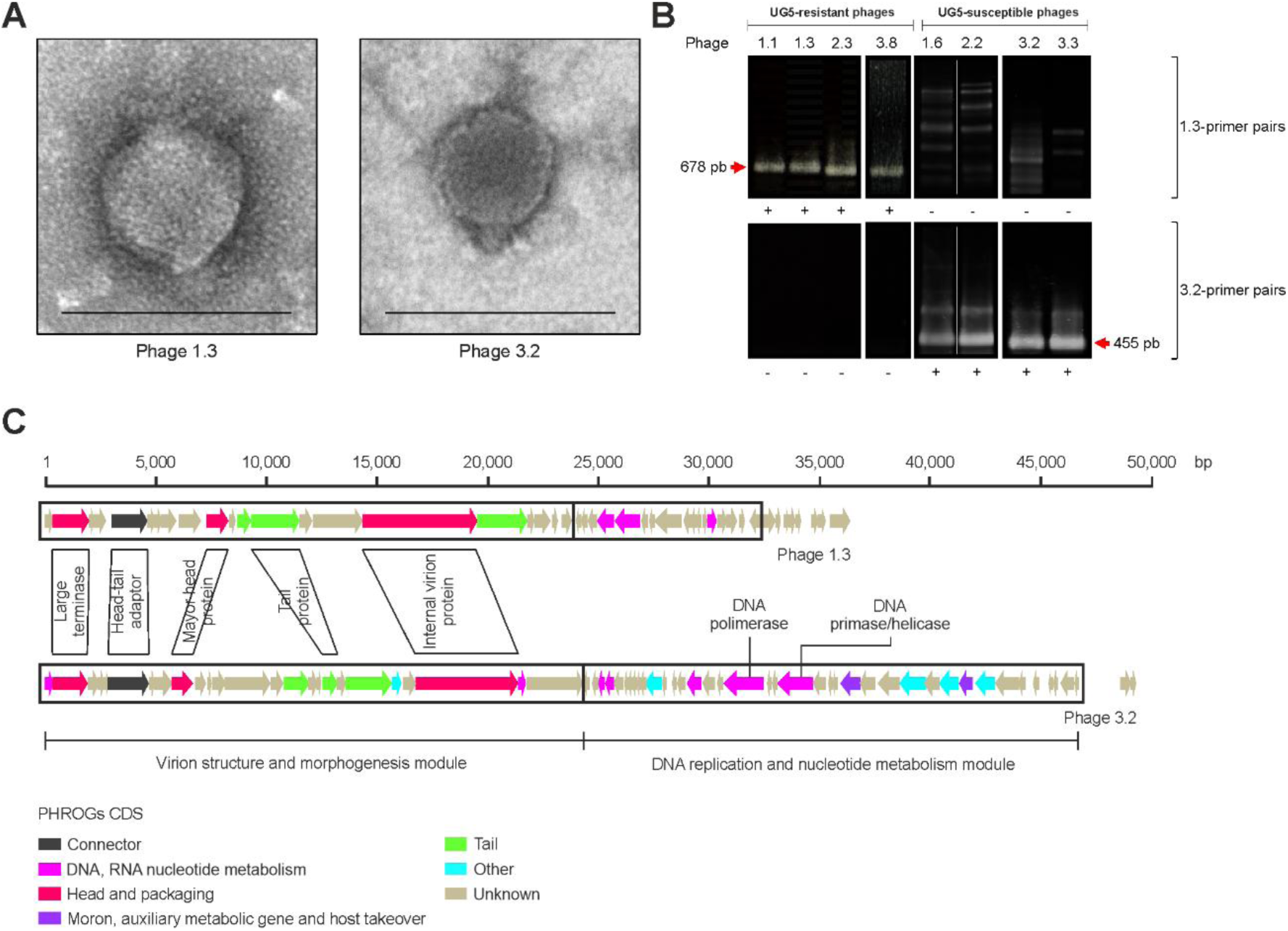
Characterization of GR4 phages. **(A)** Electron micrographs of *S. meliloti* phages 1.3 (resistant) and 3.2 (susceptible). Phage particles were negatively stained with uranyl acetate. Scale bars, 100 nm. (**B**) PCR amplification of the *terL* gene using phage-specific primer sets delineates the two phage groups that differ in their infection outcome in *S. meliloti* UG5. Phages susceptible to UG5 (represented by phage 3.2) yielded amplicons with the 3.2-specific *terL* primers, whereas phages not inhibited by UG5 (represented by phage 1.3) amplified only with the 1.3-specific primers. For simplicity, only representative phages from each group are shown. **(C)** Comparative genomic organization of phages 1.3 and 3.2. Genomic coordinates are shown in base pairs. Coding sequences (CDSs) were annotated using Pharokka (Bouras *et al*., 2023) and color-coded according to their predicted function based on PHROG (Terzian *et al*., 2021) classification: ‘connector’ (black), ‘DNA/RNA nucleotide metabolism’ (light purple), ‘head and packaging’ (red), ‘moron/auxiliary metabolic gene/host takeover’ (dark purple), ‘tail’ (green), ‘other’ (light blue), and ‘unknown’ (beige). Key structural and replication-related genes are labeled. Shading denotes homologous CDSs. Two major functional modules, virion structure and morphogenesis, and DNA replication and nucleotide metabolism, are indicated.

Given these similarities, we next asked whether phages exhibiting resistant or susceptible phenotypes to DRT11 could be distinguished by specific genomic markers. PCR amplification of the *terL* gene was therefore performed using primer pairs designed uniquely for the *terL* sequences of phages 1.3 and 3.2. Amplification with the 1.3-specific primers occurred in phages 1.1–1.3, 1.5, 1.7, 1.8, 1.9–1.11, 2.3 and 3.8, whereas the 3.2-specific primers amplified phages 1.4, 1.6, 2.1, 2.2 and 2.4–3.7 (**Figure 5B, Table S1**). This analysis delineates two clearly separated phage groups whose *terL* signatures correlate with the phenotypes defined by phages 1.3 (resistant) and 3.2 (susceptible).

To explore whether morphological similarity reflected underlying genomic relatedness, the genomes of phages 1.3 and 3.2 were sequenced and assembled using Geneious Prime (version 2024.0.7). Phage 1.3 has a genome size of 35,969 bp, whereas phage 3.2 is larger, with 48,938 bp (both under Sequence Read Archive submission SUB13566626 within BioProject PRJNA990552). Viral coding sequences (CDSs) were annotated using Pharokka (Bouras et al., 2023), and functional categories were assigned on the basis of PHROG profiles (Terzian et al., 2021), classifying genes into “connector”, “DNA/RNA nucleotide metabolism”, “head and packaging”, “moron/auxiliary metabolic gene/host takeover”, “tail”, “other”, and “unknown”. Despite differences in genome size, comparative genomic analysis revealed a conserved overall architecture in both phages, organized into two major functional modules: one associated with virion structure and morphogenesis, and another with DNA replication and nucleotide metabolism (**Figure 5C**).

## Discussion

We characterized a previously unexamined UG5-large system from *S. meliloti* RMO17, classified as a class 3 UG/Abi RT system, which confers anti-phage immunity despite lacking the transmembrane (TM) protein often associated with UG5-large loci. Our results provide experimental confirmation that TM genes are not required for UG5-large activity, in contrast to other nitrilase-associated systems such as UG1 where TM deletion compromises defense (Gao et al., 2020). We further show that a minimal two-component architecture, consisting only of the RT and its C-terminal nitrilase domain, is sufficient for antiviral function.

UG5-large modules resemble UG1 (type-I DRT) systems, such as the one described in *Klebsiella pneumoniae* NCTC9143 (Gao et al., 2020), where both the RT and nitrilase domains are required for antiviral activity. Our phylogenetic analysis of 728 nitrilase domains shows that UG1-, UG5-large-, UG5-small (CL249)-, UG6 (CL285)- and UG29-associated nitrilases fall within a single, well-supported evolutionary cluster. UG1 sequences are nested within a broader UG5-associated radiation, indicating that UG1 does not constitute an independent lineage under the present sampling, whereas UG29 forms a clearly distinct and robustly supported clade. Rather than reflecting discrete clade-specific trajectories, this topology suggests that diversification of UG-associated nitrilases has occurred within a broadly shared evolutionary space, with multiple UG modules drawing on related nitrilase lineages.

This broader phylogenetic context also clarifies the placement of a previously characterized antiviral system from *Escherichia coli* 101 (Rousset et al., 2022). Although not annotated as UG5, this system encodes an RT–nitrilase fusion that clusters within the UG5 radiation and is followed by an ORF predicted to encode a transmembrane protein, an architecture consistent with canonical UG5-large modules. Notably, the *E. coli* 101 system confers resistance to a Podoviridae phage, further supporting its functional alignment with DRT11-derived antiviral mechanisms.

The genomic positioning of DRT11 in *S. meliloti* RMO17 further supports its defensive function. PADLOC analysis shows that the locus resides within a defense-enriched region of pSymA, flanked by a nucleotidyl-transferase toxin module and two type II toxin–antitoxin systems. Although DRT11 is found in only a small subset of *S. meliloti* genomes sequenced to date, the presence of a conserved genomic scaffold in strains such as RCAM1750 and KH46 suggests that this region constitutes a hotspot for defense gene acquisition in rhizobia. Moreover, its phylogenetic proximity to a UG5-large system from *S. medicae* indicates that similar RT–nitrilase architectures may have been independently mobilized across related plant-associated species.

Functional assays in *S. meliloti* GR4 lacking DRT11 systems demonstrated that DRT11 provides robust protection against a subset of phages isolated from *Medicago sativa* rhizosphere, and mutational analysis confirmed that both catalytic domains are essential for defense. The growth phenotypes, protection at low but not high multiplicity of infection, are consistent with an abortive-infection–like mechanism and position DRT11 within the broader UG/Abi functional framework proposed by Mestre et al. (2022).

Comparative analyses of representative phages 1.3 and 3.2 revealed Podovirus-like morphology and a conserved modular genome organization, alongside marked divergence within their replication modules, a hallmark of the modular evolution characteristic of Podoviridae (Cubo et al., 2020). Although additional work is needed to identify the molecular basis of sensitivity, the separation of GR4 phages into two *terL*-defined lineages that mirror their resistant versus susceptible phenotypes suggests that DRT11 acts on a lineage-specific or replication-associated feature. Notably, despite extensive escape-mutation attempts, no DRT11-escaping phages were recovered, raising the possibility that the system targets an essential or highly constrained step in the infection cycle for which viable escape mutations are unlikely to occur.

Although the metagenomic analysis presented here was not intended to infer ecological enrichment, it indicates that UG5-associated systems are recurrently detected, albeit at low abundance, in soil- and rhizosphere-derived datasets. This observation provides a contextual framework for interpreting the functional characterization of DRT11 in a plant-associated bacterium. From an ecological standpoint, the activity of DRT11 is particularly notable given the environments in which *S. meliloti* persists. Rhizobia encounter continuous phage pressure in the bulk soil and rhizosphere, settings where viral predation influences strain persistence, competitive fitness, and ultimately the efficiency of symbiosis. That DRT11 provides robust protection against a subset of naturally occurring phages, and does so through a minimal RT–nitrilase architecture, underscores the relevance of UG-derived defenses in shaping survival strategies of plant-associated bacteria. The inability to recover escape mutants further suggests that systems like DRT11 may target infection steps that are both essential and evolutionarily constrained, a property likely to enhance their ecological impact.

As phage-based approaches expand into agricultural applications, understanding the antiviral capacities of beneficial soil bacteria becomes increasingly important for safeguarding microbial inoculants and maintaining productive plant–microbe partnerships. By defining DRT11 as a streamlined RT–nitrilase antiviral module, this work provides a foundation for exploring how reverse transcriptase–linked defenses contribute to the stability, resilience, and evolutionary dynamics of plant-associated microbiomes.

## Acknowledgments

We thank Ascensión Martos Tejera and José María del Arco Martín for technical assistance. We also thank Alicia Rodríguez-Sánchez and Andreina C. Peralta from the Confocal and Transmission Electron Microscopy Service (CTEM) at the Estación Experimental del Zaidín (CSIC) for support and technical assistance for electron microscopy EEZ service.

## Author contributions

E.S.N. performed the experimental work and contributed to writing, reviewing, and editing the manuscript. F.M.A. supervised the study and the experimental work and contributed to manuscript writing. V-M. performed RT–nitrilase mutagenesis, phage infection assays, and MOI-dependent growth experiments. M.D.M.-S. generated and curated the transcriptomic dataset. F.M.G.-R. initiated the studies on the UG5 system in *S. meliloti* RMO17. N.T.G. supervised the study, conducted the structural, metgenomic and phylogenetic analyses, and contributed to writing, reviewing, and editing the manuscript.

## Data availability

Transcriptomic data corresponding to the UG5 (DRT11) region and flanking sequences of *S. meliloti* RMO17 have been deposited in Zenodo and are available to the editors and reviewers through a private access link. The dataset will be made publicly accessible upon acceptance of the manuscript. Other data generated in this study are available from the corresponding author upon reasonable request.

## Supplementary material

### Supplementary Figures

**Figure S1.**
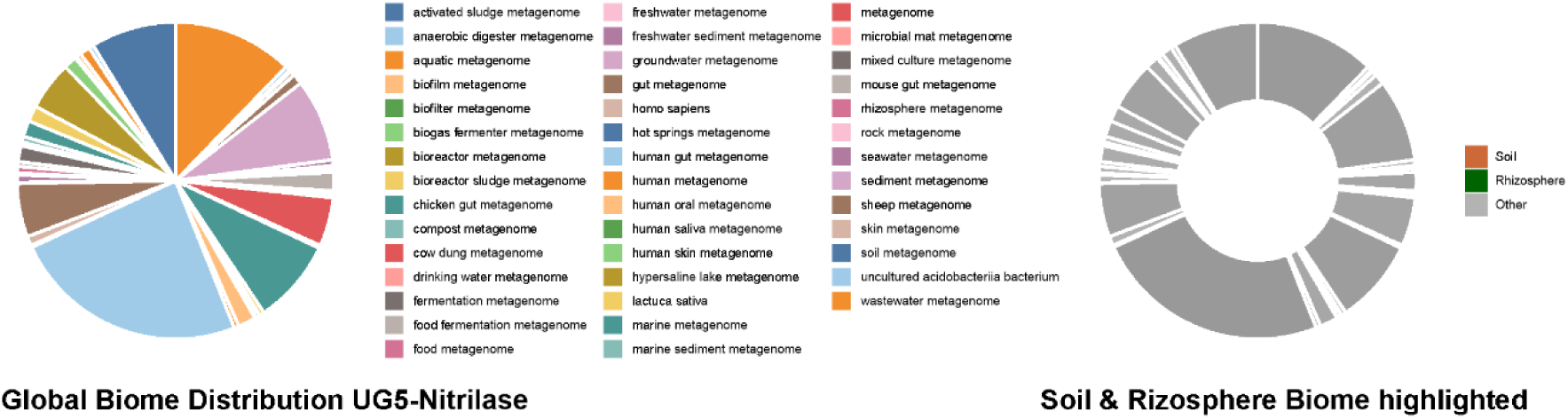
Metagenomic distribution of UG5-associated nitrilase domains across environmental biomes. UG5-associated nitrilase-domain sequences were identified by HMM-based searches against the MGnify protein database using a profile derived from the UG5 nitrilase domain. An initial set of 2,497 hits was retrieved and subsequently filtered using a stringent significance threshold (E-value ≤ 1 × 10⁻⁵⁰), yielding 1,004 high-confidence sequences. These filtered hits were then mapped to sample-level biome annotations. The figure shows the global distribution of UG5-associated nitrilase-domain sequences across annotated biomes, together with a highlighted hybrid donut representation in which soil- and rhizosphere-derived metagenomes are shown explicitly and all remaining biomes are grouped as “Other.”

**Figure S2.**
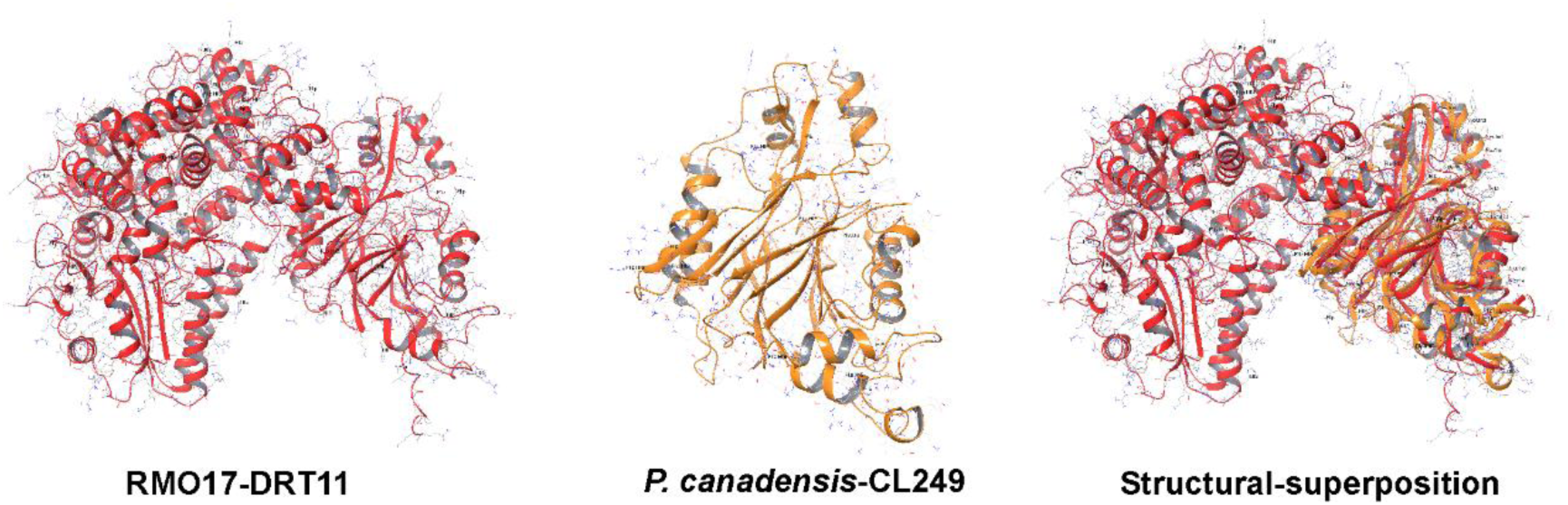
US-align structural superposition of Structure_1 (red) onto Structure_2 (orange). Structure_1 corresponds to chain A of RMO17-DRT11 (1005 residues), and Structure_2 to chain A of *Pseudomonas canadensis* (WP_14805088.1) belonging to nitrilase CL249 (307 residues). US-align produced an aligned region of 290 residues with an RMSD of 1.98 Å and a sequence identity of 0.262 across aligned positions. TM-score was 0.28047 when normalized by the length of Structure_1 (reference; L=1005, d0=10.56) and 0.88721 when normalized by the length of Structure_2 (L=307, d0=6.43).

### Supplementary Tables

**Table S1.**
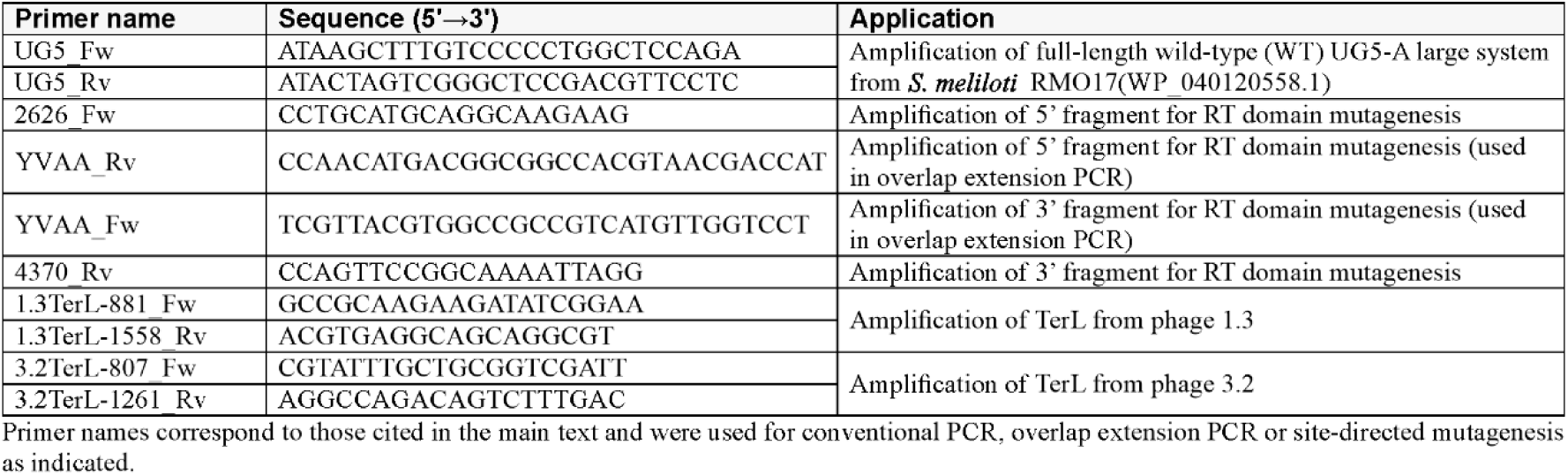
Oligonucleotides used in this study.

**Table S2.** Plasmids constructed in this study.

| Name | Description |
| --- | --- |
| pSparkI_UG5_WT | Expression of the full-length wild-type (WT) UG5-large system in the pSpark® I vector |
| pBB2_UG5_WT | Expression of the WT UG5-large system in the pBB2 vector |
| pBB2_UG5_Δ179 | Expression of the UG5-large system lacking a 179 bp region between two SmaI sites |
| pBB2.1_UG5_WT | Expression of the WT UG5-large system in the pBB2.1 vector |
| pBB2.1_UG5_RT[DD296–297AA] | Expression of the UG5-A large system carrying the DD296–297AA substitution in the RT domain in the pBB2.1 vector |
| pBB2.1_UG5_Nitrilase[C876A] | Expression of the UG5-large system carrying the C876A substitution in the nitrilase domain in the pBB2.1 vector |

All constructs are derived from the UG5-large system of *S. meliloti* RMO17 (WP\_040120558.1)

### Supplementary data

**Supplementary Data 1**. UFBoot consensus tree (.contree; Newick) obtained with IQ-TREE2 for the nitrilase-domain phylogeny in **Figure 1A**.

